# An NO-binding Cache domain receptor interacts with a Ser/Thr kinase through a conserved HAMP domain interaction

**DOI:** 10.64898/2026.05.25.727664

**Authors:** Maithili Deshpande, Yajie Xu, Robert Dunleavy, Igor B. Zhulin, Brian R. Crane

## Abstract

Cache_heme domains are a family of bacterial heme c proteins that combine a conserved Cache fold with an α-helical insertion containing a heme-binding CXXCH motif. We characterize an unusual Cache_heme–HAMP (CHH) receptor from *Pseudomonas azotoformans* with a periplasmic NO-binding Cache_heme domain, transmembrane region, and a cytoplasmic HAMP domain uncoupled from intrinsic enzymatic output. Genomic analysis reveals that CHH receptors are frequently co-localized with genes encoding Ser/Thr kinases (STPKs) that possess a catalytic domain similar to PknB, but lack peptidoglycan-binding PASTA domains. Using biochemical assays, biolayer interferometry, SEC-MALS, and cryo-electron microscopy, we show that the *P. azotoformans* CHH receptor binds its operon-associated kinase with nanomolar affinity via conserved C-terminal repeat modules in the kinase composed of two pseudo-symmetric α-helical bundles. Cryo-EM structures demonstrate that this interaction orders the otherwise flexible HAMP domain, with the repeated helical domains from the kinase contacting each HAMP subunit symmetrically. Kinase–receptor binding is ATP-independent, but kinase phosphorylation of the receptor at a specific HAMP threonine residue is substantially enhanced when NO binds to the heme c sensor domain. Thus, NO-induced conformational changes that transverse the membrane either modulate substrate accessibility in the HAMP or alter kinase activity directly. These findings potentially define a mode of bacterial NO signaling in which a periplasmic heme c receptor couples ligand sensing to cytoplasmic phosphorylation via a physically associated Ser/Thr kinase, a mechanism distinct from that of cytoplasmic NO sensors. The conserved nature of the kinase recognition motif suggests broader relevance of CHH–STPK interactions across proteobacteria.

## Introduction

We recently defined the domain of unknown function (DUF) 3365 protein family as Cache_heme, which is widely distributed across bacterial phyla, with strong enrichment in Gracilicutes^1^. Cache_heme proteins consist of a conserved Cache fold with a unique α-helical insertion containing a CXXCH heme c binding motif. Bioinformatics analysis has shown that Cache_heme proteins sequester into two families that represent either single-domain periplasmic proteins or as fusion proteins in which the Cache_heme domain is coupled to other domains likely involved in signal transduction, such as guanylate cyclase or histidine kinase domains^1^. Thus, Cache_heme proteins participate in either redox functions (single-domain proteins) or signal transduction functions (fusion proteins).

In this previous work^1^, we identified an orphan Cache_heme receptor from *P. azotoformans* that has nitric oxide (NO) binding properties and UV-Vis absorption spectra characteristic of known NO sensors. This protein has an unusual domain architecture consisting of a periplasmic Cache_heme, transmembrane (TM) helices and a cytoplasmic HAMP domain. This architecture is atypical, because HAMP domains almost always transduce conformational signals between other domains that are coupled to either their N- or C-termini^2–4^. Cache_heme proteins with only transmembrane and HAMP domains are widely distributed among γ-proteobacteria, particularly in *Pseudomonas* spp., and in β-proteobacteria such as *Burkholderia* and *Telluria* spp. (NCBI BLAST and CDART). Understanding the properties and function of this putative NO sensing receptor and its potential signaling partners will help to uncover the role that this signaling system plays in bacteria.

NO signaling is important for controlling several metabolic processes in bacteria and eukaryotes^5–8^. NO is a membrane-permeable gas molecule that is toxic at higher concentrations but serves as an important signaling molecule at lower concentration^9,10^. In eukaryotes, NO is important for vasodilation, smooth-muscle relaxation and immune response^11,12^. In bacteria, NO is involved in processes such as quorum sensing, biofilm formation and symbiosis. NO signaling typically occurs through reversible binding of NO to ferrous heme in hemoproteins, leading to conformational changes in the heme-sensing domain that are transmitted to a functional domain. The most studied heme NO sensor proteins in bacteria include the H-NOX^13–15^ (**H**eme **N**itric oxide/**O**xygen binding domain), and the more recently discovered NosP^16,17^ proteins. H-NOX proteins were discovered from sequence comparisons with eukaryotic sGCs and NosP proteins were discovered in bacteria that lack H-NOX domains. Both are cytoplasmic proteins that contain b-type hemes and participate in NO-mediated biofilm formation. The *P. azotoformans* receptor is different from these proteins in that the sensor domain is periplasmic and contains a c-type heme, which is normally matured in the periplasm. Some transmembrane heme c proteins such as *Geobacter sulfurreducens* GSU0582 and GSU0935 contain methyl-accepting chemotaxis (MCP) domains, with a periplasmic domain that has been shown to bind NO^18,19^. However, their function is still unknown, although it is believed that they may act as redox sensors.

In addition to its unusual Cache_heme domain, the *P. azotoformans* receptor lacks an enzymatic output domain and ends instead with a C-terminal cytoplasmic HAMP domain. HAMP domains are named for the proteins in which they were first discovered: **H**istidine kinases, **A**denylate cyclases, **M**ethyl-accepting chemotaxis proteins, and **P**hophatases^4^. HAMP domains are usually located between sensory input domains (typically a periplasmic ligand-sensing domain) and output (often kinase or methyl-accepting domains) and transmit conformational changes across a protein, thus playing important roles in transmembrane signaling^3,20,21^. HAMP domains (∼50 residues) are comprised of two α-helices that dimerize to form a parallel four-helix bundle^2,4,22,23^. Several models have been proposed to describe the conformational changes that the HAMP domain undergoes during signal transduction, including, 1) a swinging piston model^24,25^, 2) a gearbox model^22,26^, and 3) a dynamic model of four-helix bundle stability^27^. As the *P. azotoformans* receptor contains a HAMP domain that does not couple to an output domain, how it may transduce signals is intriguing.

In *P. azotoformans*, and in other bacterial species that contain such Cache_heme_HAMP (CHH) homologous, a Ser/Thr kinase (STPK) locates to the same operon, thereby suggesting that the receptor may regulate the kinase. In this work, we show that the receptor biochemically and structurally interacts with the kinase through its HAMP domain. The two proteins interact with nanomolar affinity, independent of ATP, through a C-terminal recognition module conserved across this subfamily of Ser/Thr kinases. Furthermore, the activity of the kinase is modulated by NO binding to the sensor domain, indicating a potential role for NO signaling through periplasmic heme c domains in bacteria.

## Results

### CHH receptors genetically associate with a predicted Ser/Thr kinase

As the *P. azotoformans* CHH receptor contains an “uncoupled” HAMP domain, its enzymatic output is likely another protein. We analyzed the genome environment of CHH receptors with the same domain architecture across bacterial species using NCBI GenBank, NCBI BLAST and the TREND (**TR**ee-based **E**xploration of **N**eighborhoods and **D**omains) platform^28^ **(Fig. S1)**. A predicted cytoplasmic Ser/Thr kinase is located in the gene operon in the majority of cases, and in addition, this kinase is nearly always present either directly upstream or downstream of the receptor. We also identified the gene for a predicted PP2C phosphatase that clusters with the receptor in some bacterial species, however, this gene is present with less frequency than that of the kinase. In the *P. azotoformans* S4 strain, the genes encoding for the kinase and phosphatase are located directly downstream of the receptor.

Conserved domain search on NCBI GenBank and NCBI BLAST revealed that the first 267 amino acid residues of the *P. azotoformans* kinase are highly conserved with those comprising the catalytic core of the Ser/Thr kinase PknB **(Fig. S2)**. PknB-type proteins are manganese (Mn)-dependent kinases that have been well-characterized in *M. tuberculosis*, *S. aureus* and *B. subtilis*^29–33^. PknB kinases have an N-terminal cytoplasmic kinase domain, transmembrane helices and multiple extracellular PASTA domains. PknB is important for a variety of functions, including peptidoglycan synthesis, cell wall metabolism, virulence, and antibiotic resistance^34–37^. However, the *P. azotoformans* kinase lacks the TM regions and PASTA domains of PknB and only shares the catalytic domain. Thus, the CHH receptor performs a different function than PknB.

AlphaFold 3 structural prediction of the Ser/Thr kinase revealed that is has an N-terminal PknB-like Ser/Thr kinase catalytic domain attached to a C-terminal domain of unknown function **(Fig. 1)**. The predicted C-terminal region consists of a ∼30 residue linker that joins to two related short four-helix bundles, also connected by a ∼30 residue linker. AlphaFold 3 predicted that the kinase and receptor interact with each other through the C-terminal domains; however, AlphaFold 3 also predicted a non-physiological interaction between the cytoplasmic catalytic kinase domain and the periplasmic receptor Cache_heme domain **(Fig. 1)**.

**Fig. 1:**
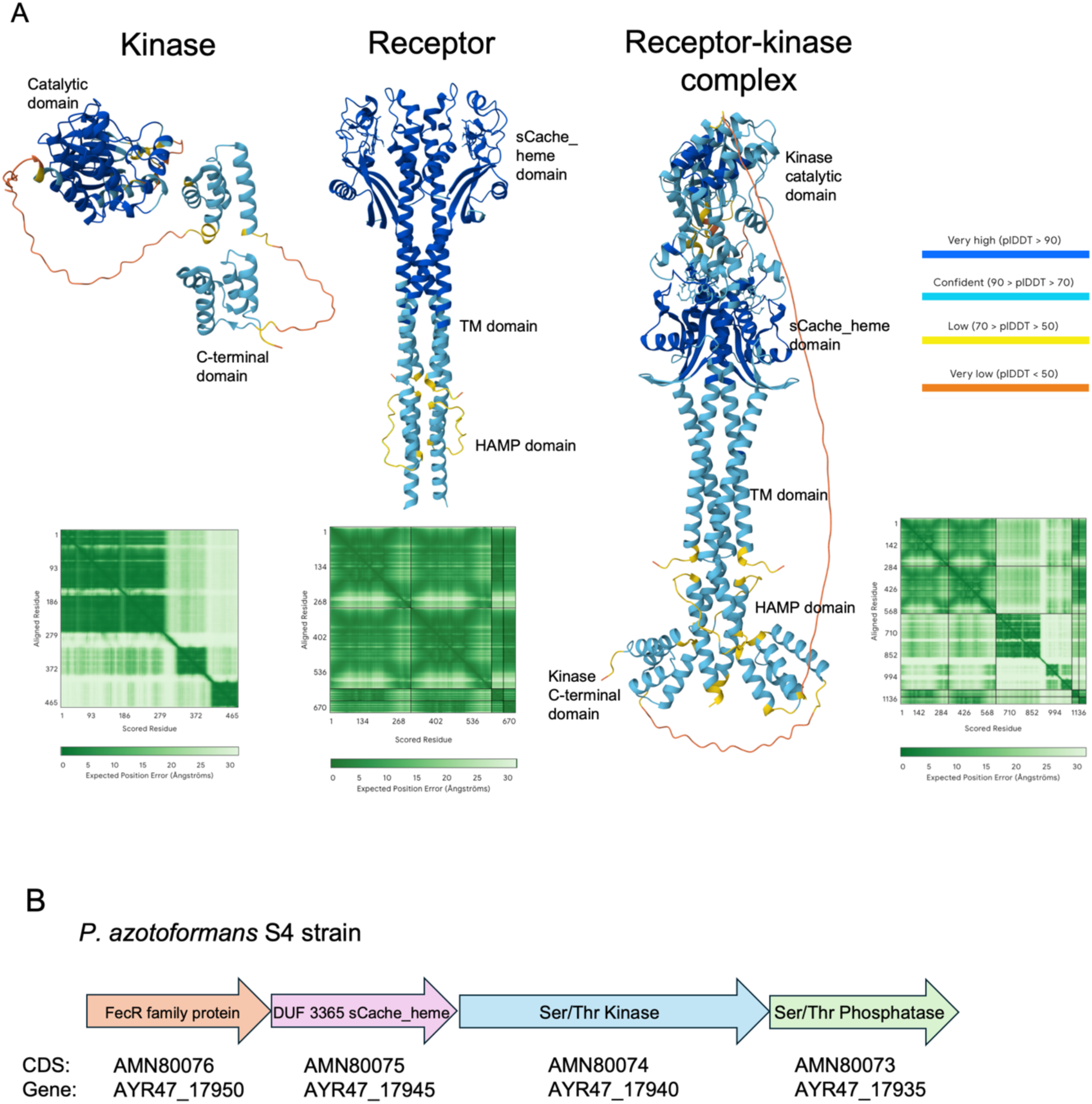
AlphaFold 3 models and gene organization of the Ser/Thr kinase, DUF 3365 Cache_heme receptor, and receptor-kinase complex. (A) The AlphaFold 3 predicted structure of the kinase consists of an N-terminal core catalytic domain and a long C-terminal domain consisting of two short helical bundles. The model of the predicted *P. azotoformans* kinase and receptor complex shows that the interaction region occurs between the HAMP domain of the receptor and the C-terminus of the kinase. AlphaFold 3 incorrectly attaches the kinase catalytic domain to the periplasmic Cache_heme domain of the receptor. (B) A conserved Ser/Thr kinase and phosphatase are present in the gene cluster of the *P. azotoformans* DUF 3365 receptor.

### Purification of the *P. azotoformans* kinase and CHH receptor

Two variants of the *P. azotoformans* kinase were expressed in *E. coli* and purified: (i) the catalytic domain (CD) comprising of the first 275 amino acid residues of the protein (STK_CD_) and (ii) the full-length (FL) kinase (STK_FL_) that includes the C-terminal domains **(Fig. S3)**. Both protein variants were expressed in *E. coli* BL21 DE3 cells with either an N-terminal 6x His tag or twin-Strep purification tag, in both soluble form and inclusion bodies. However, common *E. coli* contaminant proteins ArnA and DnaK (identified by mass spectrometry) were also highly expressed. During purification of the soluble fraction, these contaminant proteins bound with high affinity to Ni-NTA resin during purification of the His-tagged variants and thus lowered the kinase yields significantly. Additionally, these contaminants could not be separated by size-exclusion chromatography (SEC) from the FL His-tagged kinase variant because of the similar molecular weight and size of these proteins. To circumvent this problem, FL His-tagged kinase was purified by slow refolding under denaturing conditions from inclusion bodies. The His-tagged FL kinase variant was found to be active through radioisotope assays and was used in pulldown experiments with the receptor.

The twin-Strep tagged kinase variants were purified from the soluble fraction. The use of twin-Strep resin eliminated the issue of the contaminant proteins as they did not bind to Strep-resin. This procedure allowed for soluble protein purification and resulted in good protein yield. Twin-Strep tagged kinase variants were used for all biochemical experiments apart from the pulldown assays, and in structure determination of the receptor-kinase complex by cryo-electron microcopy (cryo-EM).

The *P. azotoformans* receptor was purified with an N-terminal twin-Strep tag **(Fig. S3)**. This variant was better behaved than the His-tagged form during the affinity purification step, because with the latter, the addition of imidazole during elution from Ni-NTA resin led to aggregation. The purified twin-Strep tagged form of the receptor runs as two bands of near-identical molecular size on an SDS-PAGE gel due to a slight N-terminal truncation. An anti-Strep tag western blot confirmed that the lower molecular weight band corresponds to receptor that had lost the 2.9 kDa twin-Strep tag due to N-terminal proteolysis **(Fig. S4)**. Subsequent radioisotope assays and biolayer interferometry (BLI) experiments confirmed that this small N-terminal degradation does not affect the properties of the receptor.

## The receptor and kinase interact with nanomolar affinity

We employed several different methods to test whether the receptor and kinase interact with each other and to identify which regions of the kinase are important for their interaction.

### Column affinity assays using purified receptor as bait and kinase as prey

Initial column affinity (i.e. pulldown) experiments were performed on purified receptor and His-tagged FL kinase by immobilizing the receptor on Strep-Tactin XT 4Flow resin as bait for the kinase. Kinase was flowed over resin saturated with receptor, the resin was washed and the proteins eluted with D-biotin. His-tagged STK_FL_ run over empty resin with no receptor bound was used as a control. The SDS-PAGE gels showed that STK_FL_ was pulled down by the receptor **(Fig. 2)**, but that the results for STK_CD_ were ambiguous, because it and the receptor run at approximately the same position in an SDS-PAGE gel **(Fig. S5)**.

**Fig. 2:**
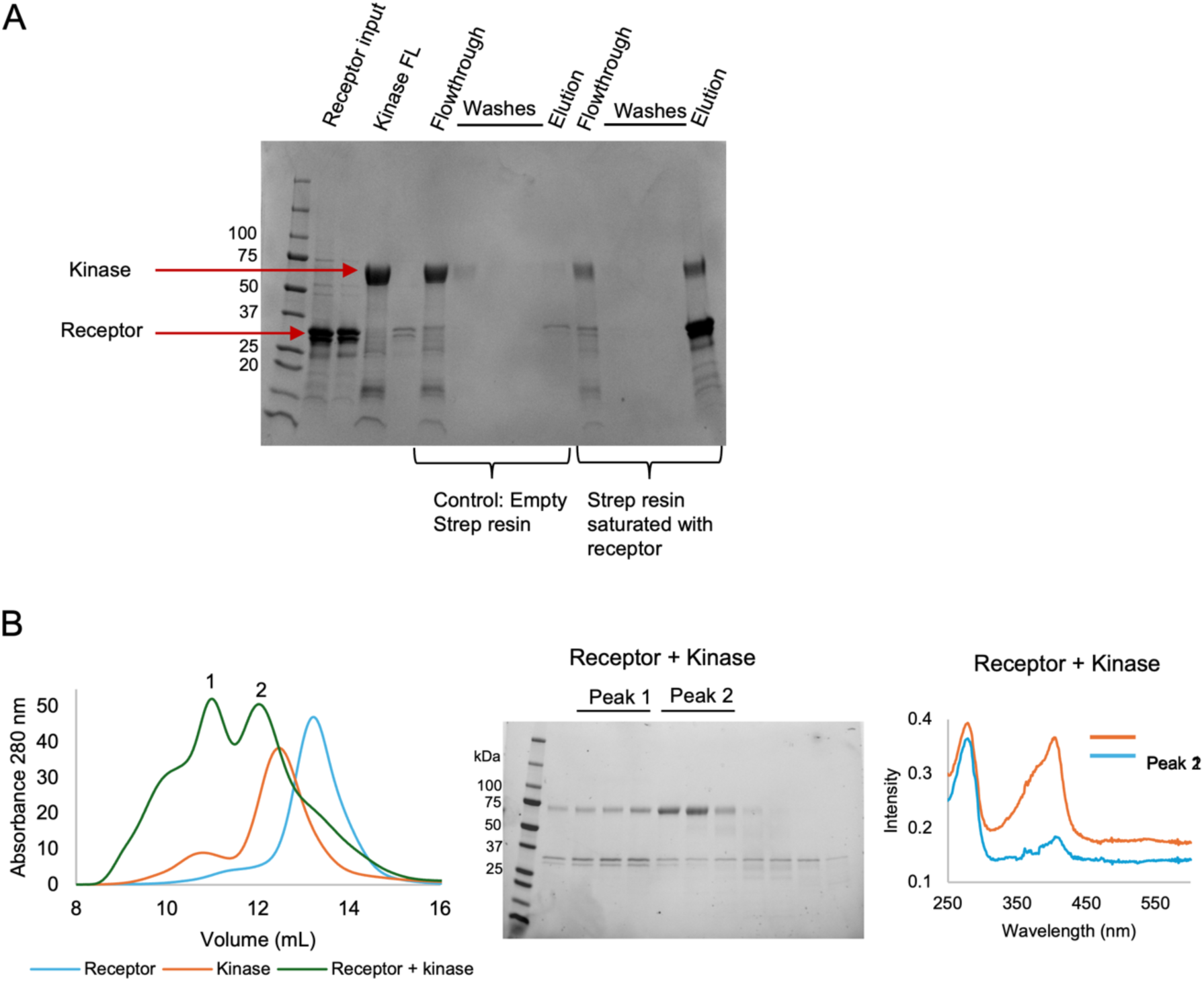
Pulldown and co-elution of the receptor-kinase complex. (A) Pulldown with immobilized receptor on resin as bait and kinase as prey. The empty resin control confirms that kinase does not bind non-specifically to resin. The receptor and kinase complex eluted together from the resin, indicating the formation of a complex. (B) The receptor-kinase complex co-elutes from the size-exclusion chromatography (SEC) column in peak 1 (left). SDS-PAGE gel of fractions obtained from SEC of the receptor and kinase solution (middle). UV-Vis absorption spectra on peaks 1 and 2 (right). The receptor-kinase mixture eluted earlier from the column in two peaks. UV-Vis spectroscopy and SDS-PAGE gels run on the fractions showed that peak 1 (10.5 mL) corresponds to a receptor-kinase complex whereas peak 2 (12.5 mL) mainly consists of unbound kinase.

### Co-elution of the receptor and kinase using size-exclusion chromatography (SEC)

An equimolar ratio of STK_FL_ to CHH receptor monomer was co-eluted on SEC at an earlier fraction compared to receptor or kinase alone (**Fig. 2**). Though peak 1 and 2 on the chromatogram have similar amount of protein as seen by absorbance at 280 nm in the UV-Vis spectra, the Soret peak at 407 nm is much higher for peak 1 than peak 2, indicating that there is low amount of receptor in peak 2. This indicates that the majority of the receptor elutes earlier in a receptor-kinase mixture (10.5 mL) than it does without kinase present (13.5 mL). Thus, the receptor and kinase co-elute as a complex from an SEC column. The high amount of unbound kinase and low amount of unbound receptor in the complex also suggests that the stoichiometric ratio of complex formation is one kinase molecule per receptor dimer. On incubating receptor and kinase at a ratio of 1 kinase:1 receptor dimer, most of the kinase complexes with the receptor **(Fig. S6)**. Again, coelutions could not be used to test interaction between the truncated kinase and receptor due to the smaller difference in their molecular weights.

Size-exclusion chromatography coupled to multi-angled light scattering (SEC-MALS) was also performed on the receptor, kinase and receptor-kinase complex **(Fig. S7)**. The receptor eluted as two peaks, likely due to degradation of the receptor as SEC-MALS was performed at room temperature and requires high protein concentration. The first major peak of 119 kDa matches the molar mass estimated for dimeric protein with the LMNG detergent micelle and agrees with the estimated molecular weight obtained for the His-tagged variant of the receptor from small-angle x-ray scattering (SAXS) experiments^1^. The molar mass of the kinase obtained from MALS was overestimated as ∼65 kDa, compared to the ∼55 kDa molecular weight calculated from the amino acid sequence. This is likely due to the flexible C-terminal domains and the presence of detergent micelles in the buffer causing the protein to run differently on the SEC column that a typical globular protein. For the receptor-kinase mixture, we used a stoichiometric ratio of 1 kinase molecule per receptor monomer. The first elution peak of the receptor-kinase incubated mixture has a molecular weight of ∼190 kDa, which agrees with the sum of the individual molecular weights of receptor and kinase obtained by MALS.

### Bio-layer interferometry (BLI)

BLI was used to measure binding kinetics of STK_FL_ and STK_CD_ to the receptor. Experiments were performed in the presence and absence of ATP to determine if ATP-binding to the kinase had an effect on the interaction. We immobilized biotinylated FL or truncated kinase on Octet Streptavidin (SA) biosensor tips and the association and subsequent dissociation of a dilution series of the receptor to immobilized kinase was recorded. The dissociation constant (K_d_) for receptor interaction with STK_FL_ calculated by steady-state analysis was ∼90 nM **(Fig. 3)**. We observed no difference in the K_d_ in the presence of ATP and Mn^2+^, thus the kinase domain does not have to be bind cofactors for interaction with the receptor. There was no binding between the receptor and truncated kinase regardless of the presence or absence of ATP and Mn^2+^, thus corroborating the AlphaFold 3 model that the C-terminal domains are required for receptor-kinase complex formation.

**Fig. 3:**
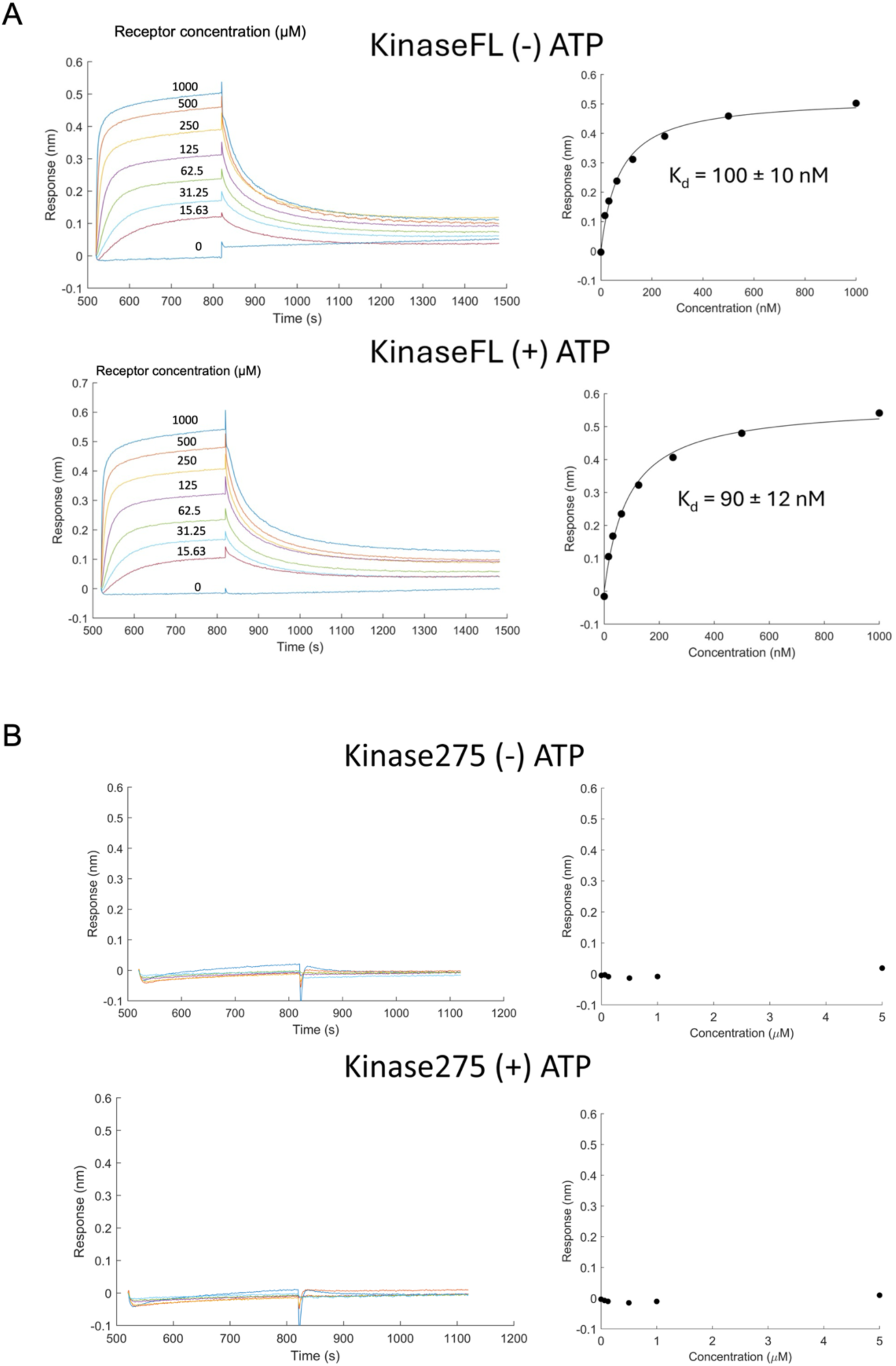
Biolayer interferometry (BLI) of receptor to STK_FL_ binding kinetics (A) and STK_CD_ lacking the C-terminal domain (B) in the presence and absence of ATP. (A) BLI traces of the association and dissociation of a series of receptor concentrations on a biosensor tip loaded with FL kinase (left). Steady-state analysis (right) reveals K_d_ values of 100 nM and 90 nM respectively in the presence and absence of ATP and Mn^2+^. The affinity does not depend upon ATP-binding to the kinase. (B) BLI traces of receptor association and dissociation to truncated kinase (left) and steady-state analysis (right). The receptor and kinase do not interact non-transiently regardless of ATP binding to the truncated kinase.

## Ligands bound to the receptor influence kinase activity in the complex

### Phosphorylation of the receptor by the kinase increases when receptor is bound to NO

The amino acid sequence and structural fold of the catalytic domain of the *P. azotoformans* kinase were predicted to be similar to those of the catalytic domain of Mn^2+^-dependent kinase PknB. We therefore initially tested the auto-phosphorylation activity of STK_FL_ and STK_CD_ with radio-labeled γ^32^-P-ATP under buffer conditions containing different concentrations of MnCl_2_ and MgCl_2_ **(Fig. S8)**. Similar to PknB, Mn^2+^ is required for catalytic activity and cannot be substituted by Mg^2+^.

To test if the receptor had a regulatory effect on modulating kinase activity, we incubated three states of the receptor – reduced ferrous heme (Fe(II)), oxidized ferric (Fe(III)) and ferrous NO-bound anaerobically with an equimolar amount of kinase prior to performing phosphorylation assays **(Fig. 4)**. Whereas the kinase phosphorylated the receptor in all three states, the rate of phosphorylation of the NO-bound receptor is significantly higher than reduced or oxidized states. This lends support to our hypothesis based on the UV-Vis absorption spectra and NO binding kinetics^1^ that the receptor may play a role in NO-sensing. The increase in receptor phosphorylation suggests that the receptor undergoes a conformational change on binding NO, either exposing additional phosphorylation sites, or making the same phosphorylation site more accessible to the kinase.

**Fig. 4:**
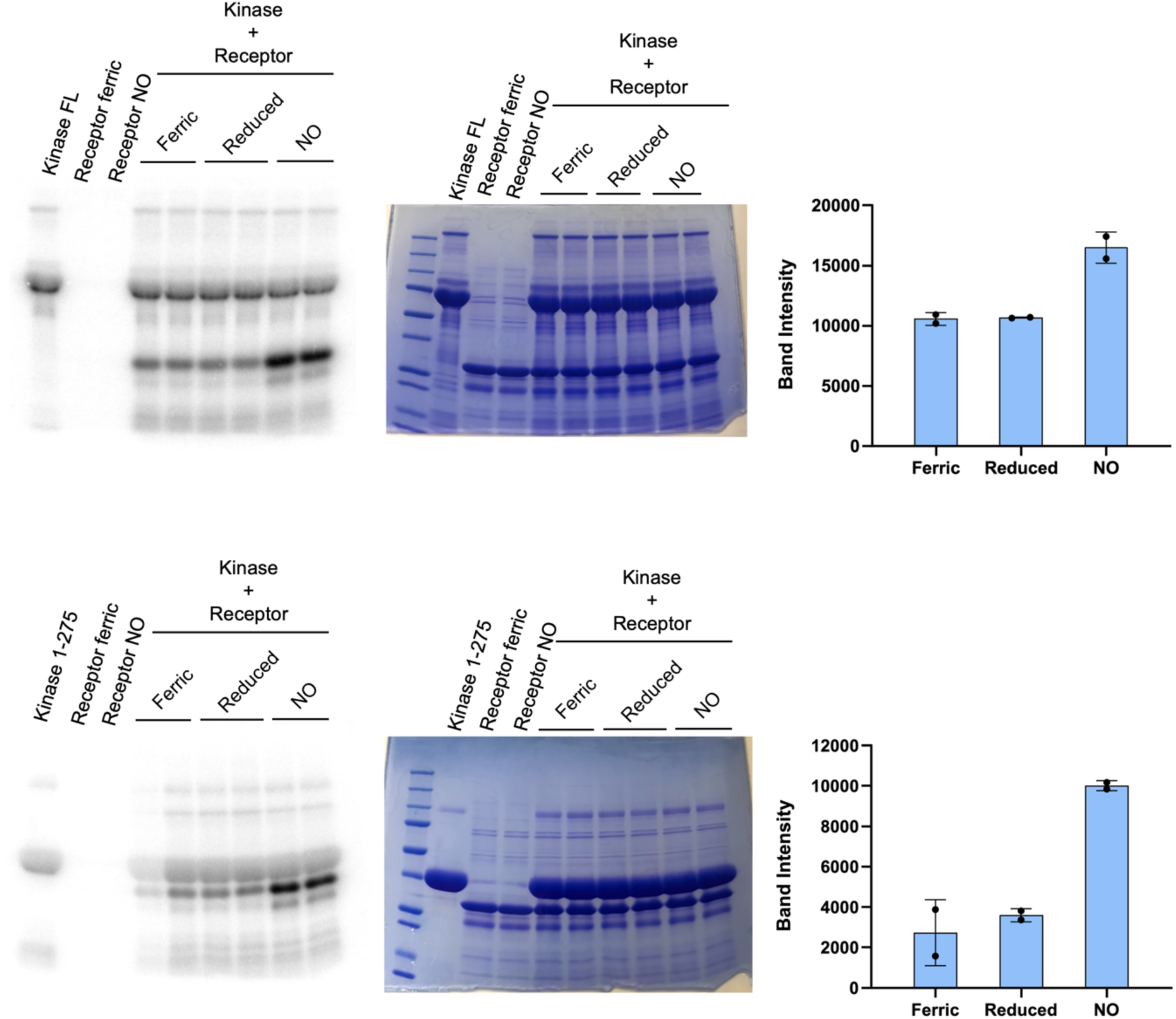
Autoradiograms of receptor phosphorylation in ferric, ferrous and NO-bound states by STK_FL_ (A) and STK_CD_ (B) (A) Phosphorylation of receptor by STK_FL_. The autoradiogram (left) shows that NO-bound ferrous receptor undergoes more phosphorylation than the ferric or reduced (ferrous) state. The Coomassie-stained gel (middle) confirms that protein loading in all wells is the same. Band intensities were quantified by ImageJ (right). (B) Phosphorylation of receptor by STK_CD_ with autoradiogram (left), Coomassie-stained gel (middle) and band intensity plot (right). A similar trend is observed in that the kinase phosphorylates the NO-bound receptor at higher rates.

### Characterization of the phosphorylation sites by mass spectrometry

To identify phosphorylated S/T sites in the receptor, we performed the phosphorylation reaction using equimolar amounts of the receptor and either SKT_FL_ or SKT_CD_ with excess ATP **(Fig. S9)**. Residue Thr256 in the HAMP domain was phosphorylated by both kinase variants. This residue is located at the start if a flexible loop linking the two helices of the HAMP domain in each subunit. Importantly, the enhanced phosphorylation occurs at the cytoplasmic HAMP domain, where the kinase is localized, but on the other side of the membrane from the NO-binding heme c domain.

## The receptor-kinase complex forms with NO-bound and reduced states of the receptor

As the state of the heme moiety affects receptor phosphorylation, we performed pulldown assays anaerobically with the reduced ferrous form of the CHH receptor and the NO-bound state **(Fig. S10)**. Similar affinity binding experiments as described above were performed with ferrous and NO-bound receptor immobilized on Strep-Tactin resin under anaerobic conditions. The FL kinase bound to the receptor in both cases. Thus, heme reduction and NO binding does not alter receptor-kinase complex formation to an appreciable extent.

## Cryo-electron microscopy of the free CHH receptor and the receptor-kinase complex

To further investigate receptor–kinase interactions, we performed single-particle cryo-EM analysis on specimens prepared from LMNG-solubilized receptor alone and in complex with the kinase (**Table S1**).

3,396 dose-fractionated movies were collected on a 200 keV Talos Arctica for the receptor only specimen with 149,071 particles selected for the 3D refinement (**Fig. S10**). A 5.07 Å structure of the isolated *P. azotoformans* receptor reveals a protein dimer with clearly discernible periplasmic heme-containing Cache domain and transmembrane helices within the detergent micelle **(Fig. 5)**. However, little to no density was obtained for the HAMP domain. This lack of structural resolution is likely because the unbound HAMP domain is highly flexible and is also relatively small compared to the bulky detergent micelle above it.

**Fig. 5:**
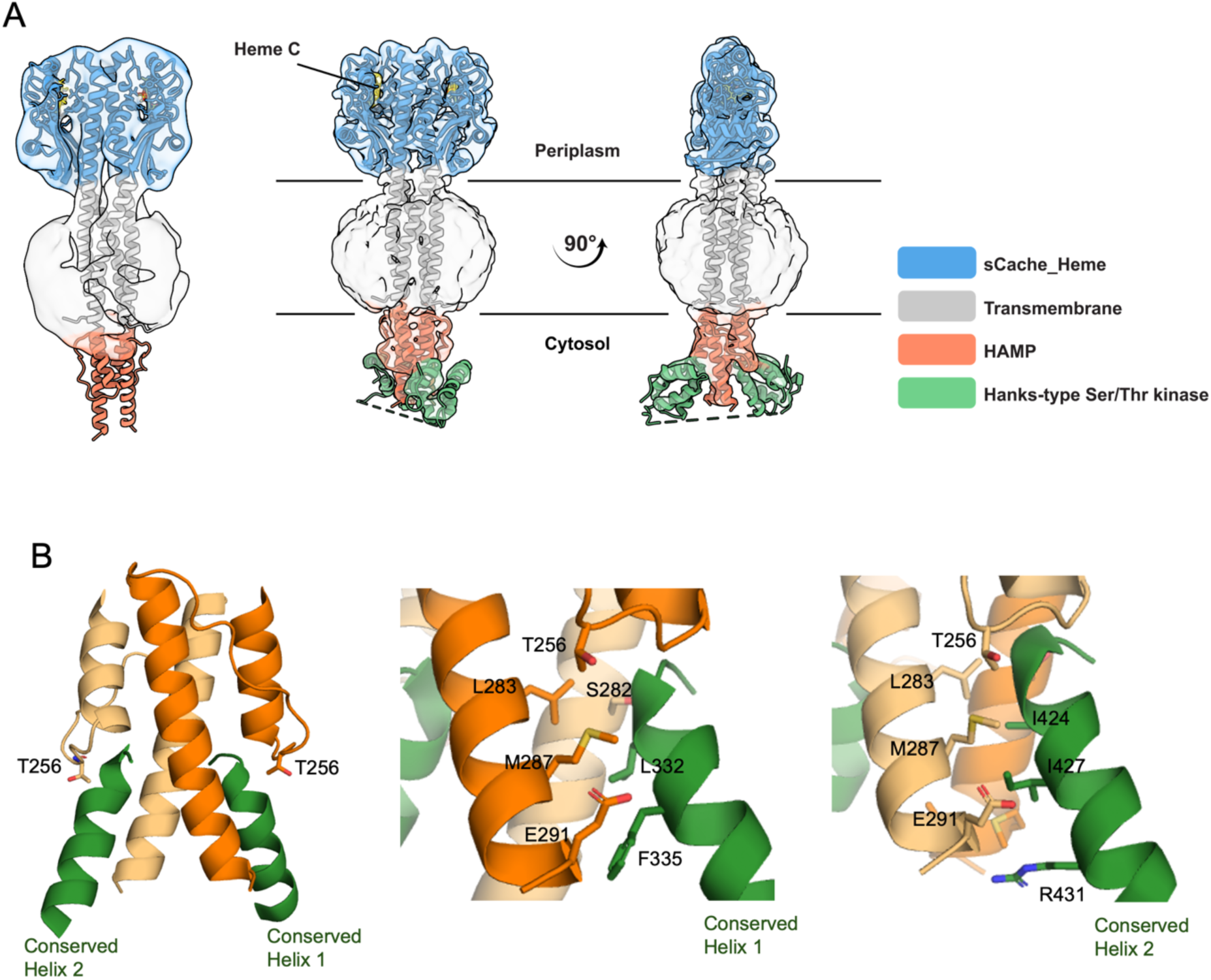
Cryo-EM structures of the receptor and receptor-kinase complex. (A) Density map of receptor aligned with the AlphaFold 3 model (left). Transmembrane helices are clearly apparent within the detergent micelle and the extracellular domain conforms to that expected for the Cache_heme fold. Density map of receptor-kinase complex aligned with the AlphaFold 3 model (right). The HAMP domain is well discerned in the structure along with the interaction region of the kinase. The C-terminal domain of the kinase interacts with the HAMP domain of the receptor. Electron density for the catalytic domain of the kinase was not visible. (B) Zoomed view of the interaction region of the receptor and kinase. Two short highly conserved residue sequences conserved among kinases of this type align beneath HAMP helix AS-1 to interact with the receptor (green).

∼15,800 dose-fractionated movies were collected on a 300 keV Titan Krios for the receptor-kinase complex and 179,400 particles were kept for final refinements from the selected 2D classes (**Fig. S11**). This data led to a 3.47 Å resolution density map of the complex that reveals that the kinase C-terminal domain binds to and orders the HAMP domain and provides a detailed picture of the interaction between receptor and kinase. The kinase interacts with the HAMP domain of the receptor pseudo-symmetrically through the two C-terminal α-helical bundles. In each of the two helical domains, related sequences IGPLARFLVRKSLN (domain 1, conserved helix 1) and IGPIARIVVSRALR (domain 2, conserved helix 2), form α-helices that closely associate with the two symmetric subunits of the HAMP domain (**Fig. 5**). Sequence searchers performed on the recognition consensus sequence reveal that it is highly conserved as a dyad repeat in Ser/Thr kinases of this type throughout gamma proteobacteria. Thus, the kinase C-terminal helical domains likely function as protein recognition motifs for similar HAMP domains in CHH receptors.

## Discussion

The *P. azotoformans* DUF 3365 receptor has an unusual domain architecture consisting of a periplasmic heme c Cache_heme domain (previously DUF 3365), a transmembrane region and an isolated cytoplasmic HAMP domain (hence a “CHH” receptor). An output “uncoupled” HAMP domain is uncommon; HAMP domains nearly always transmit signals or conformational changes between a sensor and enzymatic output module in the same protein. We have shown that the *P. azotoformas* CHH receptor physically associates and modulates the activity of a genetically linked STK kinase as its output model. Remarkably, NO-binding to the ferrous form of the receptor alters kinase-mediated phosphorylation of the HAMP domain, to which the kinase binds with highly conserved small helical binding modules. Furthermore, BLI data confirmed that the receptor and kinase interact with high affinity and similar K_d_ values in the presence and absence of ATP and Mn^2+^.

Previously^1^, we have shown that the *P. azotoformans* receptor behaves similarly to known b-type heme NO sensor proteins such as NosP and sGC in terms of UV-Vis spectral features and NO dissociation kinetics. Here, we have shown that the kinase phosphorylates the receptor at a higher rate when NO is bound to the receptor, in an Mn^2+^ dependent manner. This activation by NO further suggests that the receptor and its interacting partners represents a coupled system for NO signaling in bacteria. We do allow that another, yet unknown, heme ligand could also activate the system. Nevertheless, the effect of NO does demonstrate that ligands to the heme moiety of the sCache_heme domain can transmit conformational signals across the membrane that alter the conformation of the HAMP is such a way that its ability to act as a substrate for the bound kinase changes. Given how the kinase binds the HAMP, through the non-catalytic C-terminal binding modules that are coupled flexibly to the SKT_CD_, it seems unlikely that the receptor would modulate the activity of the catalytic domain directly.

The majority of NO-binding proteins are cytoplasmic and contain non-covalently bound heme b. One of the few examples of heme c periplasmic proteins having strong NO affinity is cytochrome c’ in proteobacteria^38–40^. NO binding to these proteins induces physiological effects and they have been shown to scavenge externally supplied NO^41,42^. The *P. azotoformans* receptor has similarities to group 1 cytochromes c’ in both UV-Vis spectra and in the solvent accessibility to the heme moiety^38^.

A periplasmic heme c receptor that is responsive to NO begs the question of why gram-negative bacteria would evolve such a signaling system? A sensing domain in the periplasm would generate a potentially faster response as the NO would not have to cross the cytoplasmic membrane and may also lead to a more localized response with less interference or dilution of the signal by other processes in the cytoplasm that react with NO. Oxygen is another potential ligand for a type c heme, but we have shown previously that O_2_ does not bind stability to the Cache_heme and rather rapidly autoxidizes the cofactor^1^. Here we show that heme oxidation/reduction alone does not alter kinase activity to the extent that NO does. We cannot rule out some other effector of the system, but given the ability of NO to propagate conformational change to the HAMP, if there is another ligand, it likely coordinates the heme. Whereas we do not currently know the further downstream signaling consequences of this particular system, we note that NO sensors in bacteria are often involved in NO-mediated biofilm formation, quorum sensing and metabolism^7,8,43^.

Bacterial Ser/Thr kinases play roles in cell signaling, metabolism, virulence and secretory processes, among others^44^. The catalytic domain of the kinase located in the CHH operon contains the standard residues important for phosphotransferase activity, namely the ATP-binding lysine (here, VALK Lys), catalytic aspartate (HRD Asp) and the metal-binding Asp (DFG Asp)^45,46^. Bacterial STPKs often have additional C-terminal domains specific for function. For example, PknB in *Mycobacterium tuberculosis* and *Streptococcus pneumoniae* contain extracellular PASTA domains that bind peptidoglycans to activate the STKm which then phosphorylates its substrates and starts a signaling cascade to regulate cell division and peptidoglycan synthesis. *Yersinia pestis* YpkA, important for virulence, contains a C-terminal guanine nucleotide inhibitor (GDI) domain and an actin-binding domain^47,48^. Unlike the more studied STKs in bacteria, the kinase in the CHH protein contains two ∼50 psuedo-symmteric a-helical bundles. These C-terminal domains directly recognize the HAMP domain through conserved ∼15 residue sequences These two kinase binding motifs are highly conserved among STKs of this type, hence associating them with HAMP modules in their cognate receptors.

## Materials and Methods

### Cloning

The coding sequence for codon-optimized full-length *Pseudomonas azotoformans* kinase (GenBank: AMN80074) was synthesized by Twist Bioscience in vector pET8a with an N-terminal His tag for expression in *E. coli*. The truncated kinase consisting of the first 275 residues (STK_CD_) was cloned by Kinase-Ligase and Dpn1 (KLD) cloning. Constructs of the *P. azotoformans* receptor, full-length and truncated kinase with an N-terminal twin Strep tag replacing the His-tag were also cloned using KLD cloning. All plasmids were confirmed by DNA sequencing at Eurofins or the Cornell University Biotechnology Center.

### Protein expression and purification

#### Purification of the twin-Strep tagged receptor with LMNG

The receptor was expressed recombinantly in *E. coli* and membrane fractions were prepared with the same method as was used for the His-tagged protein as described preivoulsy. The following differences in purifying the twin-Strep-tagged protein were employed: The supernatant obtained after solubilization and centrifugation was incubated with Strep Tactin XT 4Flow resin for 1 h at 4°C. The protein was slowly transferred into buffer containing lauryl-maltose neopentyl glycol (LMNG) by washing with a decreasing amount of buffer containing DDM, followed by washing with buffers containing LMNG. Resin was exchanged with 10 column volumes of wash buffer 1 (25 mM Tris pH 8.5, 500 mM NaCl, 10% glycerol, 0.1% DDM) followed by 5 column volumes of wash buffer 2 (25 mM Tris pH 8.5, 500 mM NaCl, 10% glycerol, 0.1% LMNG) and 10 column volumes of wash buffer 3 (25 mM Tris pH 8.5, 500 mM NaCl, 10% glycerol, 0.01% LMNG). Resin was incubated with elution buffer (25 mM Tris pH 8.5, 500 mM NaCl, 10% glycerol, 0.01% LMNG, 50 mM D-biotin) for 1 h and the protein elution was then collected. The elution was then run over an S200 16/60 gel filtration column pre-equilibrated with GF buffer (25 mM Tris 8.5, 250 mM NaCl, 5% glycerol). Fractions containing protein were pooled, protein was stored at 4°C and used within 72 h.

#### Purification of the kinase variants

The *P. azotoformans* kinase variants were expressed in *E. coli* BL21 (DE3). Cells were grown in Terrific broth (IBI Scientific) containing 50 µg/mL kanamycin at 37°C to an OD 600 of ∼0.8 and induced with 0.5 mM IPTG at 18°C overnight. Cells were harvested after ∼16 h by centrifugation at 4000 x g for 15 min at 4 °C. Cells were lysed using an Avanti Emulsiflex C3 high pressure homogenizer and then centrifuged for 1 h at 75,000 xg. The purification for the different variants then differed as follows: *STK_FL_:*

This kinase could not be purified as soluble protein because it could not be separated from common contaminants DnaK and ArnA owing to high non-specific binding to Ni-NTA resin and the similar molar mass of these proteins and the kinase. A significant amount of kinase was present in inclusion bodies, and thus this kinase was purified under denaturing conditions to remove the contaminants and then slowly refolded. Cells were lysed in lysis buffer (25 mM Tris pH 8.0, 500 mM NaCl, 10% glycerol) and the supernatant was discarded. To obtain pure inclusion bodies, the pellet was resuspended in lysis buffer supplemented with 2% Triton X-100 and centrifuged to separate membrane proteins in the supernatant. A high salt wash was then performed on the pellet using lysis buffer containing 1 M NaCl, followed by a lower salt wash. The inclusion bodies were then resuspended in lysis buffer supplemented with 6 M urea. The supernatant obtained after centrifugation was incubated with Ni-NTA resin, and resin was washed with 20 CV of the same buffer. Kinase was eluted in elution buffer (25 mM Tris pH 8.0, 500 mM NaCl, 10% glycerol, 300 mM imidazole, 6 M urea). The elution was rapidly diluted 1:1 in lysis buffer to reduce the urea content to 3 M and then slowly refolded for 12 h each in two steps by dialysis. The protein was first dialyzed into lysis buffer containing 1 M urea, and then finally into lysis buffer. Refolded protein was concentrated and then injected ono an S200 16/60 SEC column. Fractions were pooled, concentrated, frozen in liquid nitrogen and stored at -80 °C.

#### STK_CD_ His-tagged

Although high amounts of DnaK and ArnA were also present during the expression of this protein, the size and behavior of the truncated kinase was different enough that it could be purified as soluble protein and separated by SEC. After cell lysis in lysis buffer, protein was incubated with Ni-NTA resin. The resin was then washed in wash buffer (25 mM Tris pH 8.0, 500 mM NaCl, 10% glycerol, 20 mM imidazole) and eluted in elution buffer (25 mM Tris pH 8.0, 500 mM NaCl, 10% glycerol, 300 mM imidazole) prior to injection on an S200 16/60 SEC column.

#### Full-length and truncated twin-Strep tagged kinase

Supernatant after lysis and centrifugation were incubated with Strep Tactin XT 4Flow resin and then washed with 20 CV of lysis buffer and eluted in 25 mM Tris 8.0, 500 mM NaCl, 10% glycerol, 50 mM D-biotin. Elution was concentrated and injected onto the SEC column.

Column affinity assays of kinase and receptor:

Column affinity (i.e. pull-down) experiments were performed by saturating 50 μl of Strep Tactin XT 4Flow resin with twin-Strep tagged receptor and then washing off any excess. His-tagged kinase variants were passed over the resin. Resin that was not saturated with receptor was used as a control to test for non-specific binding. Resin was washed multiple times with 10 CV buffer (25 mM Tris pH 8.5, 250 mM NaCl, 5% glycerol, 0.0025% LMNG). Proteins were eluted in 25 mM Tris pH 8.5, 250 mM NaCl, 5% glycerol, 0.0025% LMNG, 50 mM D-biotin. SDS-PAGE gel samples were prepared of all flowthroughs, washes and elutions.

Pulldowns performed with ferrous and NO-bound states of receptor were done anaerobically. Ferrous and NO-bound states of receptor were prepared by using the method described previously^1^. Kinase variants used for these experiments were degassed for at least 12 h in the anaerobic chamber prior to use.

#### Size-exclusion chromatography coupled to multi-angle light scattering (SEC-MALS)

SEC-MALS experiments were carried out on free receptor, free kinase, and receptor-kinase complex. The complex was prepared by incubating an equimolar ratio of receptor and kinase for 2 h at 4 °C. For each sample, 100 μl 5 mg/ml of protein was injected onto an S200 10/300 Increase column connected to a Wyatt Technology Optilab T-rEX refractometer. Bovine serum albumin (BSA) was used as a protein standard.

#### Biolayer interferometry (BLI)

BLI experiments were performed on a Sartorius Octet RH-16 system. Full-length and truncated kinase were biotinylated by incubating with 20x molar excess of EZ-Link Sulfo-NHS-Biotin (Thermo Scientific) overnight. Protein was then desalted to remove excess and non-reacted reagent. Biotinylated kinase at a concentration of 100 nM was immobilized on Octet Streptavidin SA sensor tips. Two-fold serial dilutions of receptor were prepared in a 96-well plate with concentrations ranging from 1 μM to 15.6 nM. All proteins were in the same buffer during the experiment (25 mM HEPES 8.0, 250 mM NaCl, 5% glycerol, 0.0025% LMNG). To test if ATP-binding had an effect on the receptor-kinase, the buffer used was 25 mM HEPES 8.0, 250 mM NaCl, 5% glycerol, 0.0025% LMNG, 1 mM MgCl_2_, 1 mM MnCl_2_, 100 μM ATP. The following parameters were applied for the BLI experiment: 100 s baseline, 300 s loading (kinase), 120 s baseline, 300 s association (receptor), and 300 s of dissociation (receptor). The raw data was fit to a 2:1 heterogeneous binding model on the Octet software, and K_d_ was calculated based on steady-state analysis.

#### Radioisotope phosphorylation assays

Samples were prepared in activity buffer (25 mM Tris pH 8.5, 250 mM NaCl, 5 % glycerol, 0.0025% LMNG, 1 mM MgCl_2_, 1 mM MnCl_2_) using 10 μM of kinase and 10 μM of receptor in a 25 μl reaction volume. For ferrous and NO-bound receptor samples, the experiment was performed anaerobically. The phosphorylation reaction was initiated by the addition of 1 mM ATP mixed with radiolabeled γ-^32^P-ATP at a final radioactivity of 0.15 ± 0.05 mCi. The reaction was quenched after 10 min using Laemmli SDS-PAGE sample buffer containing 10 mM EDTA. Samples were run on a tris-glycine gel for 1 h 30 min at 110 V. The gels were stained in Coomassie, then dried and placed in a radio-cassette for 48-72 h. The phosphor screen was imaged on a Typhoon FLA-7000 imager. Band intensities were quantified using ImageJ software.

### Characterization of the phosphorylation sites by mass spectrometry

#### Sample preparation

The receptor was phosphorylated by incubating 10 μM of receptor and 10 μM of kinase in activity buffer supplemented with 5 mM ATP overnight at 4°C. The reaction was quenched in SDS PAGE Laemmli sample buffer and run on an SDS PAGE gel. The band corresponding to receptor was cut out and submitted to the Proteomics and Metabolomics Facility of Cornell University for analysis.

#### In-gel trypsin digestion of SDS gel bands

The protein band from an SDS-PAGE gel was sliced into ∼1 mm cubes and subjected to in-gel digestion followed by extraction of the tryptic peptide as reported previously^49^. The excised gel pieces were washed consecutively with 200μL distilled/deionized water followed by 50mM ammonium bicarbonate, 50% acetonitrile and finally 100% acetonitrile. The dehydrated gel pieces were reduced with 50μL of 10mM DTT in 100mM ammonium bicarbonate for 1 hour at 56 °C, alkylated with 50μL of 55mM iodoacetamide in 100mM ammonium bicarbonate at room temperature in the dark for 45 minutes. Repeated wash steps as described above. The gel was then dried and rehydrated with trypsin (Promega), at an estimated 1:10 w/w ratio in 50mM ammonium bicarbonate, 10% ACN (100ul used to overlay gel @ 10ng/ul trypsin, total enzyme used 1 ug). Incubated at 37 °C for 18 hrs. The digested peptides were extracted twice with 200μl of 50% acetonitrile, 5% formic acid and once with 200μl of 75% acetonitrile, 5% formic acid. Extractions from each sample were pooled together and filtered with 0.22 um spin filter (Costar Spin-X from Corning) and dried to dryness in the speed vacuum. Each sample was reconstituted in 0.5% formic acid prior to LC MS/MS analysis.

#### LC/MS/MS Analysis

The in-gel tryptic digests were reconstituted in 20 μL - 0.5% FA for nanoLC-ESI-MS/MS analysis, which was carried out using an Orbitrap Fusion^TM^ Tribrid^TM^ (Thermo-Fisher Scientific, San Jose, CA) mass spectrometer equipped with a nanospray Flex Ion Source, and coupled with a Dionex UltiMate3000RSLCnano system (Thermo, Sunnyvale, CA). The gel extracted peptide samples (2.5 μL) were injected onto a PepMap C-18 RP nano trapping column (5 µm, 100 µm i.d x 20 mm) at 20 µL/min flow rate for rapid sample loading and then separated on a PepMap C-18 RP nano column (2 µm, 75 µm x 25 cm) at 35 °C. The tryptic peptides were eluted in a 90 min gradient of 5% to 35% acetonitrile (ACN) in 0.1% formic acid at 300 nL/min., followed by an 8 min ramping to 90% ACN-0.1% FA and an 8 min hold at 90% ACN-0.1% FA. The column was re-equilibrated with 0.1% FA for 25 min prior to the next run. The Orbitrap Fusion is operated in positive ion mode with spray voltage set at 1.6 kV and source temperature at 275°C with advanced peak determination. External calibration for FT, IT and quadrupole mass analyzers was performed. In data-dependent acquisition (DDA) analysis, the instrument was operated using FT mass analyzer in MS scan to select precursor ions followed by 3 second “Top Speed” data-dependent CID ion trap MS/MS scans at 1.6 m/z quadrupole isolation for precursor peptides with multiple charged ions above a threshold ion count of 10,000 and normalized collision energy of 30%. MS survey scans at a resolving power of 120,000 (fwhm at m/z 200), for the mass range of m/z 375-1575. Dynamic exclusion parameters were set at 45 s of exclusion duration with ±10 ppm exclusion mass width. All data were acquired under Xcalibur 4.3 operation software (Thermo-Fisher Scientific).

#### Data analysis

The DDA raw files for CID MS/MS were subjected to database searches using Proteome Discoverer (PD) 2.5 software (Thermo Fisher Scientific, Bremen, Germany) with the Sequest HT algorithm. Processing workflow for precursor-based quantification. The PD 2.5 processing workflow containing an additional node of Minora Feature Detector for precursor ion-based quantification was used for protein identification. The database search was conducted against *E.coli* Uniprot database with added a target protein DUF 3365 receptor, which contains 15,055 sequences. Two-missed trypsin cleavage sites were allowed. The peptide precursor tolerance was set to 10 ppm and fragment ion tolerance was set to 0.6 Da. Variable modification of methionine oxidation, deamidation of asparagine/glutamine, phosphorylation of serine/threonine/tyrosine, protein N-terminal M loss, N-terminal acetylation and M-loss+acetylation, and fixed modification of cysteine carbamidomethylation, were set for the database search. To confidently localize phosphorylation sites, the phosphoRS 3.0 node integrated in PD 2.5 workflow was also used. The algorithm of phosphoRS 3.0 software enables automated and confident localization of phosphosites by determining individual probability values for each putatively phosphorylated site within the validated peptide sequences. Only high confidence peptides defined by Sequest HT with a 1% FDR by Percolator were considered for the peptide identification.

All phosphorylated peptides were validated by manual inspection of the relevant MS/MS spectra. The occupancy rate of all phosphorylated peptide was calculated based on the peak area of XICs versus their counterpart native peptides in different charges states using Xcalibur software under assumption that ionization efficiency is the same or similar between the native peptide and its phosphorylated form.

### Cryo-electron microscopy

For cryo-EM sample preparation, 4 ul of purified protein was applied to PELCO easiGLOW^TM^ plasma cleaned grids (Quantifoil R 1.2/1.3 Cu 200 Mesh and Au 300 Mesh). Grids were blotted using a Vitrobot Mark IV (FEI) set to 4 °C and 100% humidity and plunge-frozen in liquid ethane. The DUF3365 receptor only dataset was collected at a magnification of 79,000X with pixel size of 1.07 Å, on a 200 keV Talos Arctica (Thermo Fisher Scientific) equipped with a Gatan K3 detector and a BioQuantum-LS energy filter (20 eV slit width). Data were acquired at the Cornell Center for Materials Research (CCMR) using SerialEM. The data collection of DUF3365 receptor-kinase complex was performed at the National Center for Cryo-EM Access and Training (NCCAT) using Leginon (Suloway et al., 2005) on a Titan Krios (Thermo Fisher Scientific) operated at 300 kV at 105000x magnification in super-resolution mode with the calibrated pixel size of 0.4135 Å.

Those dose-fractionated movies were imported and preprocessed in cryoSPARC (v4.6.0 or later). Micrographs containing non-vitreous ice and poor CTF fits were excluded from further processing. Particles were initially picked using the blob picker and subsequently cleaned through rounds of 2D classification, ab-initio reconstructions and heterogeneous refinement. The resulting particles were then used for Topaz training within cryoSPARC following with 3D reconstructions and refinements prior to reference based motion correction (RMBC). The final density was fit to the AlphaFold 3 models of the receptor and receptor-kinase complex. Data processing workflows are shown in **Fig. S11 and S12**.

## Acknowledgements

These studies were supported by NIH grant GMR35122535. We thank the National Center for CryoEM Access and Training (NCCAT) and the Cornell Center for Material Science (CCMR) for access to cryo-EM data collection facilities.

## Competing interests

The authors declare that they have no competing interests.

Data and materials availability: The cryoEM data and structures are available at the Protein Data Bank with accession codes: 35YX and 35YV and EMDatabase maps: EMD-77289, EMD-77290. All other data needed to evaluate the conclusions in the paper are present in the paper or the Supplementary Materials.

