## Supplementary material for "An NO-binding Cache domain receptor interacts with a Ser/Thr kinase through a conserved HAMP domain interaction": Fig. S1

| Dataset | sCache_heme HAMP receptor | sCache_heme HAMP receptor Hanks-type Ser/Thr kinase |
|---|---|---|
| **State** | Homodimer | Heterotrimer |
| **Data Collection and Processing** | | |
| **Microscope** | Arctica | Krios |
| **Voltage (keV)** | 200 | 300 |
| **Detector** | K3 | Gatan K3 |
| **Nominal magnification** | 79,000 | 105,000 |
| **Data Acquisition Software** | Serial EM | Leginon |
| **Electron dose (e−/Å$^2$)** | 50 | 52.72 |
| **Pixel Size (Å) (binned )** | 1.07 | 0.4135(0.827) |
| **Defocus range (μm)** | 0.6-2 | 0.3-2.2 |
| **Number of movies (#)** | 3,396 | 7,138 +13,708 |
| **Number of particles** | 149,071 | 179,400 |
| **Symmetry imposed** | C1 | C1 |
| **Resolution (Å)** | 5.07 | 3.47 |
| **FSC threshold** | 0.143 | 0.143 |
| **Refinement** | | |
| **Initial model used** | AlphaFold3 | AlphaFold3 |
| **Atoms** | 9,288 (4670 hydrogens) | 11,575 (5825 hydrogens) |
| **Protein residues** | 584 | 728 |
| **Ligand(#)** | HEC (2) | HEC (2) |
| **RMS Deviation** | | |
| **Bond lengths (Å)** | 0.004 | 0.003 |
| **Bond angles (°)** | 0.654 | 0.66 |
| **MolProbity score** | 1.63 | 1.75 |
| **Clashscore** | 7.22 | 7.95 |
| **Poor rotamers (%)** | 2.01 | 1.61 |
| **Ramachandran Favored (%)** | 98.97 | 97.08 |
| **Ramachandran Allowed (%)** | 1.03 | 2.92 |
| **Ramachandran Disallowed** | 0.0 | 0.0 |
| **Fit to map (CCmask)** | 0.7 | 0.8 |
| **EMDB (maps)** | EMD-77290 | EMD-77289 |
| **PDB (model)** | 35YX | 35YV |

Table S1: Cryo-EM data collection and structure refinement statistics.

A

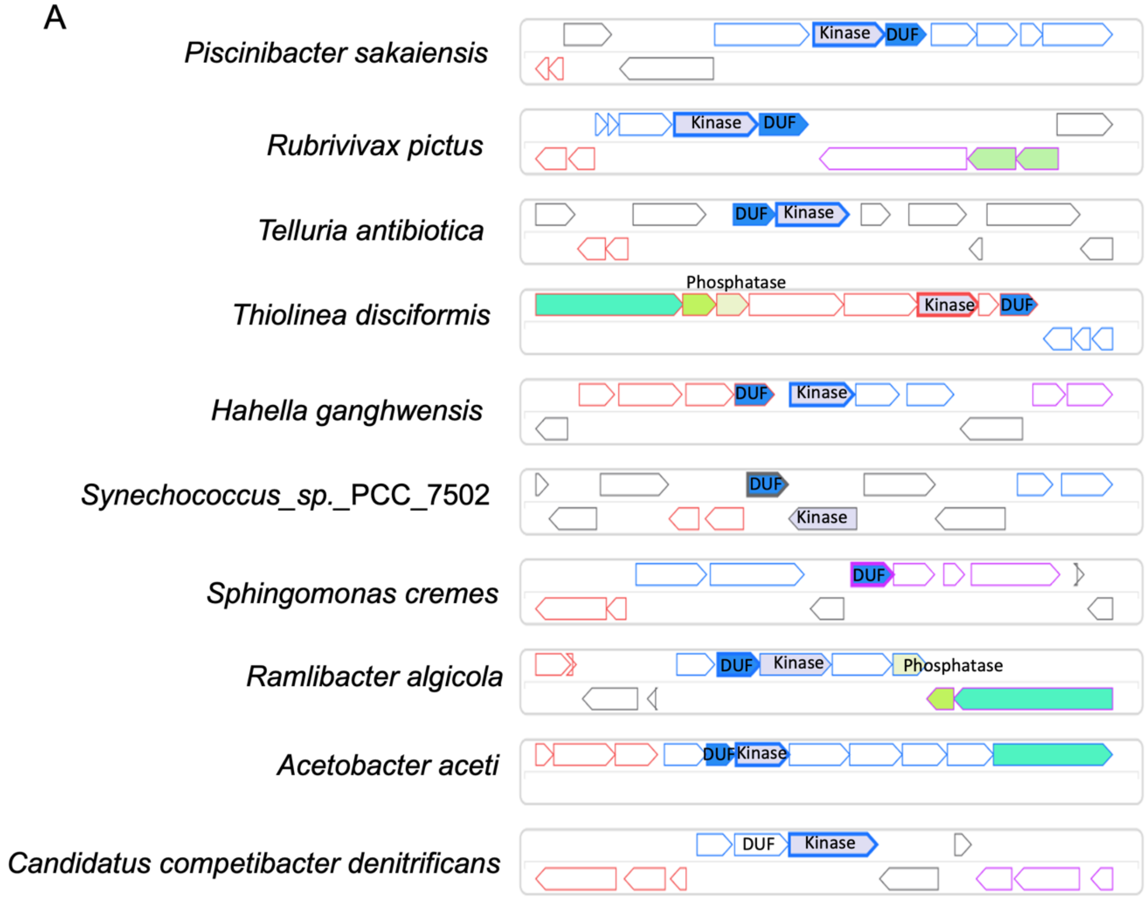

Fig. S1: Gene locus of DUF 3365 CHH receptors in selected bacterial species.

The gene neighborhoods of CHH receptors having the domain architecture of periplasmic sCache_heme domain, transmembrane (TM) region and cytoplasmic HAMP domain of selected bacterial species. The gene neighborhoods were identified using **TR**ee-based **E**xploration of **N**eighborhoods and **D**omains (TREND). The Ser/Thr kinase, DUF 3365 sCache_heme receptor, and phosphatase are annotated on the gene neighborhood diagram.

```
CLUSTAL O(1.2.4) multiple sequence alignment

P.azotoformans      ---MLEQLGKYRIDSVLGKGAMGTVYKAFDPNIARTVALKTIRKELFGDSQHAELVSRFK      57
M.tuberculosis      MTTPSHLSDRYELGEILGFGGMSEVHLARDLRLHRDVAVKVLRADLA---RDPSFYLRFR      57
S.aureus            -MIGKIINERYKIVDKLGGGGMSTVYLAEDTILNIKVAIKAIFIPPR---EKEETLKRFE      56
B. subtilis         MLIGKRISGRYQILRVIGGGGMANVYLAEDIILDREVAIKILRFDYA---NDNEFIRRFR      57
                             :*.:   :* *.*. *: * *  :   **:* :        .. .   **.

P.azotoformans      NEAQAAGRLMHPNIVAVYDYGEDE----GSAYIAMEFVEGTPLNTLLAAQTPRDLSHSLG      113
M.tuberculosis      REAQNAAALNHPAIVAVYDTGEAETPAGPLPYIVMEYVDGVTLRDIVHTEGPMTPKRAIE      117
S.aureus            REVHNSSQLSHQNIVSMIDVDEED----DCYYLVMEYIEGPTLSEYIESHGPLSVDTAIN      112
B. subtilis         REAQSASSLDHPNIVSIYDLGEED----DIYYIVMEYVEGMTLKEYITANGPLHPKEALN      113
                    .*.: :. * *  **:: * .* :       *:.**:::*  *   : :. *   . ::

P.azotoformans      WMRQLLLALEYAHSKGVVHRDIKPANLLITADNHVKVTDFGVARLDS---STLTQTGSMI      170
M.tuberculosis      VIADACQALNFSHQNGIIHRDVKPANIMISATNAVKVMDFGIARAIADSGNSVTQTAAVI      177
S.aureus            FTNQILDGIKHAHDMRIVHRDIKPQNILIDSNKTLKIFDFGIAKALS--ETSLTQTNHVL      170
B. subtilis         IMEQIVSAIAHAHQNQIVHRDIKPHNILIDHMGNIKVTDFGIATALS--STTITHTNSVL      171
                       :   .: .:*.  ::***:** *::*     :*: ***:*   :   .::*:*  ::

P.azotoformans      GTPSYMSPEQFCGELIDGRSDVFSAGIVLYQLLTGERPFSGSATMV--MQQILNQTPVAP      228
M.tuberculosis      GTAQYLSPEQARGDSVDARSDVYSLGCVLYEVLTGEPPFTGDSPVSVAYQHVREDPIPP-      236
S.aureus            GTVQYFSPEQAKGEATDECTDIYSIGIVLYEMLVGEPPFNGETAVSIAIKHIQDSVPNVT      230
B. subtilis         GSVHYLSPEQARGGLATKKSDIYALGIVLFELLTGRIPFDGESAVSIALKHLQAETPSAK      231
                    *:  *:****  *      :*::: * **:::*.*. ** *.: :    :::  .

P.azotoformans      SSLNLALDPALDSVIVRALAKRPADRYPSAHAFLDDLEALLGTTTGSWTSADIDDDRTVL      288
M.tuberculosis      SARHEGLSADLDAVVLKALAKNPENRYQTAAEMRADLVRVHNGEPPEAPKVLTDAERTS-      295
S.aureus            TDVRKDIPQSLSNVILRATEKDKANRYKTIQEMKDDLSSVLHENRANEDVYELDKMKTI-      289
B. subtilis         -RWNPSVPQSVENIILKATAKDPFHRYETAEDMEADIKTAFDADRLNEKRFTIQEDEEMT      290
                       .  :   :. ::::*  *   .** :   :  *:         .      :  .
```

Fig. S2: Sequence alignments of the catalytic domain of the *P. azotoformans* kinase and PknB-type kinases from *Mycobacterium tuberculosis*, *Staphylococcus aureus* and *Bacillus subtilis* using ClustalW.

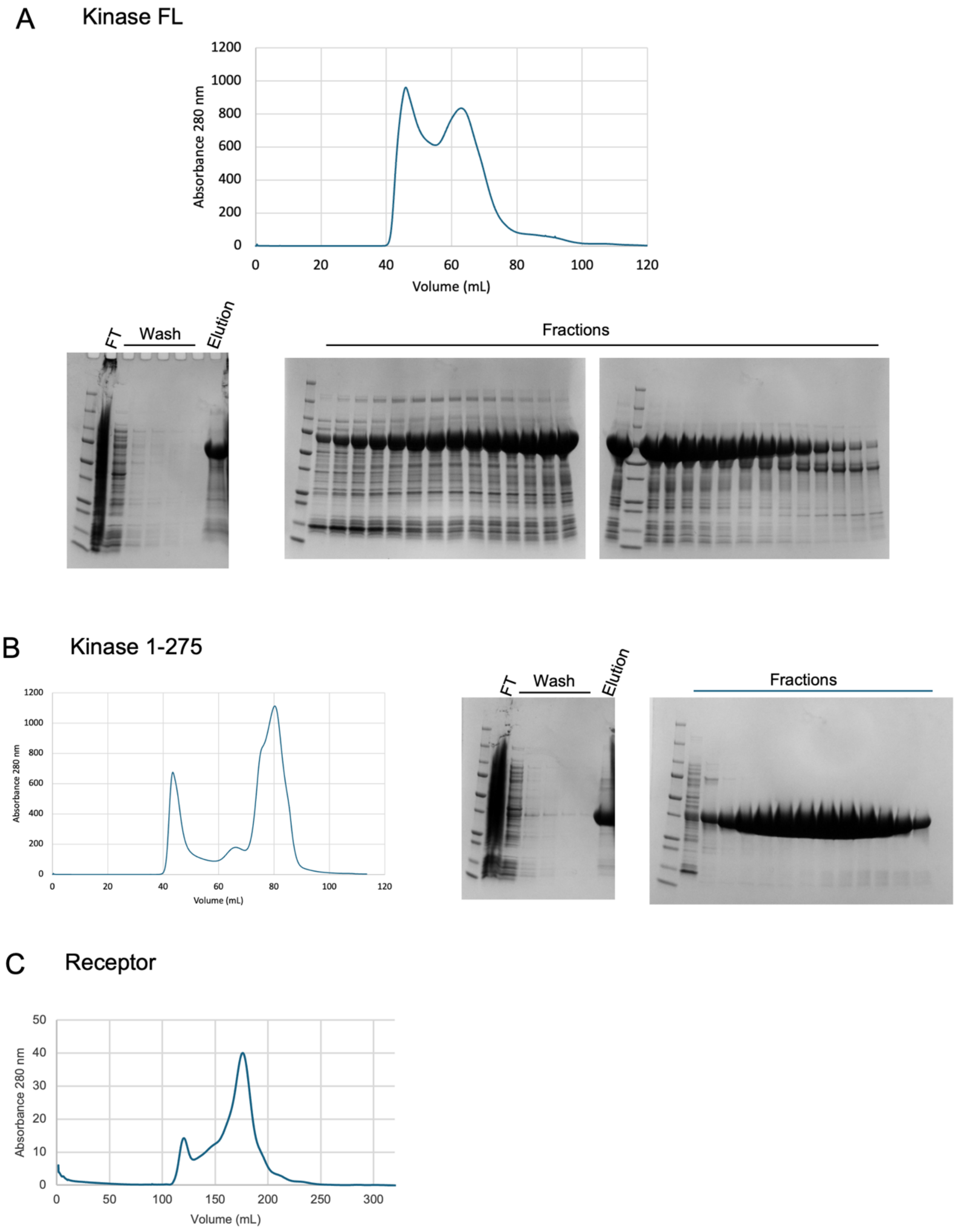

Fig. S3: Purification of twin-Strep tagged full-length kinase (A), truncated 1-275 kinase (B), and receptor (C). SEC elution profiles are shown with gel bands of associated fraction. Affinity purification gels (flow through (FT), Wash and Elution lanes) are shown to the left.

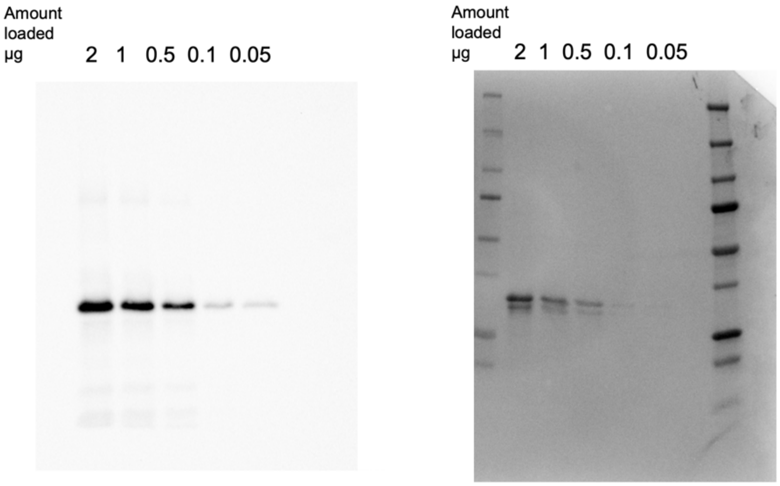

Fig. S4: Anti-Strep tag western blot on the purified twin-Strep tagged *P. azotoformans* receptor.

Comparison of the imaged blot (left) and Coomassie-stained membrane (right) indicates that the lower band in the *P. azotoformans* receptor is a slight N-terminal truncation; the lower band lacks the twin-Strep tag.

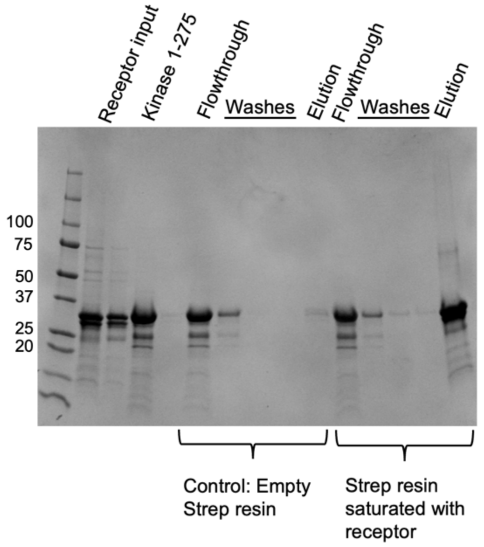

Fig. S5: Pulldown of the Ser/Thr kinase catalytic domain

The catalytic domain of the kinase and the receptor run at similar sizes in an SDS-PAGE gel. Thus, this method cannot be used to conclusively determine an interaction between $STK_{CD}$ and the receptor, although the banding pattern does indicate that the slightly largely kinase co-elutes with the receptor.

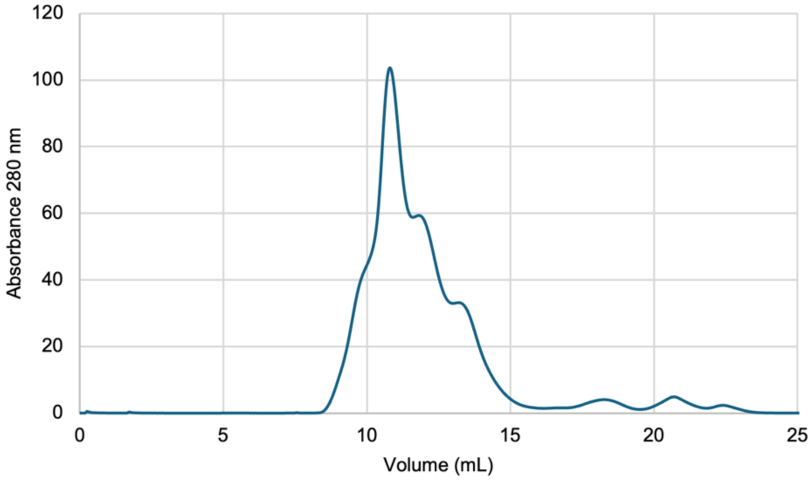

Fig. S6: Co-elution of receptor and FL kinase on SEC incubated in a (1 receptor dimer:1 kinase) ratio

Majority of the receptor complexes with the kinase when incubated together at a stoichiometric ratio of 1 dimer:1 kinase.

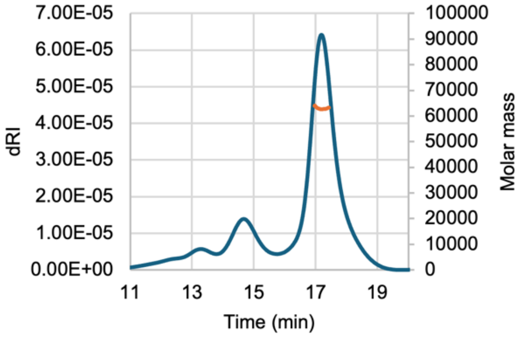

BSA MW = 63.08 ± 0.934%

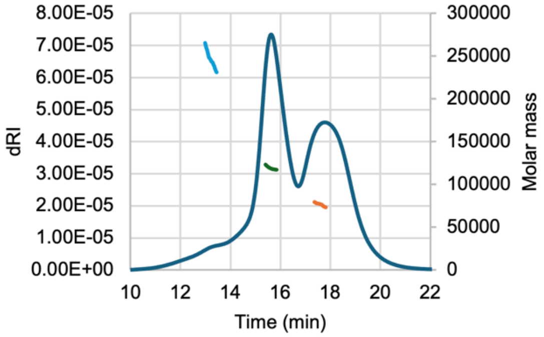

Receptor

Peak 1 = 245.8 ± 3.81%
Peak 2 = 119 ± 0.991%
Peak 3 = 76.34 ± 8.986%

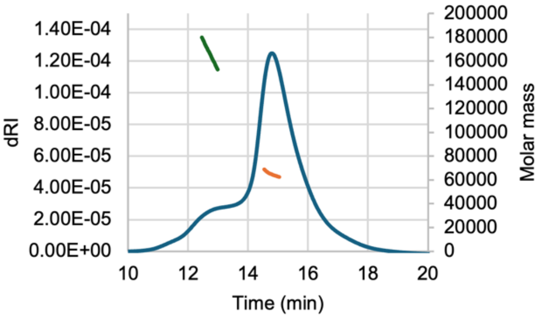

Kinase

Peak 1 = 165.7 ± 1.377%
Peak 2 = 64.99 ± 0.660%

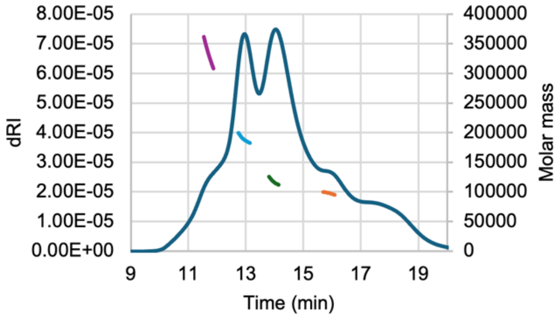

Receptor + Kinase

Peak 1 = 332.2 ± 0.872%
Peak 2 = 189.4 ± 0.658%
Peak 3 = 117.2 ± 0.828%
Peak 4 = 97.66 ± 1.617%

.

Fig. S7: SEC-MALS chromatograms with calculated molecular weights for BSA, receptor, kinase, and receptor-kinase incubation mixture (values are in kDa). The receptor-kinase complex elutes from the column as peak 2 (bottom right).

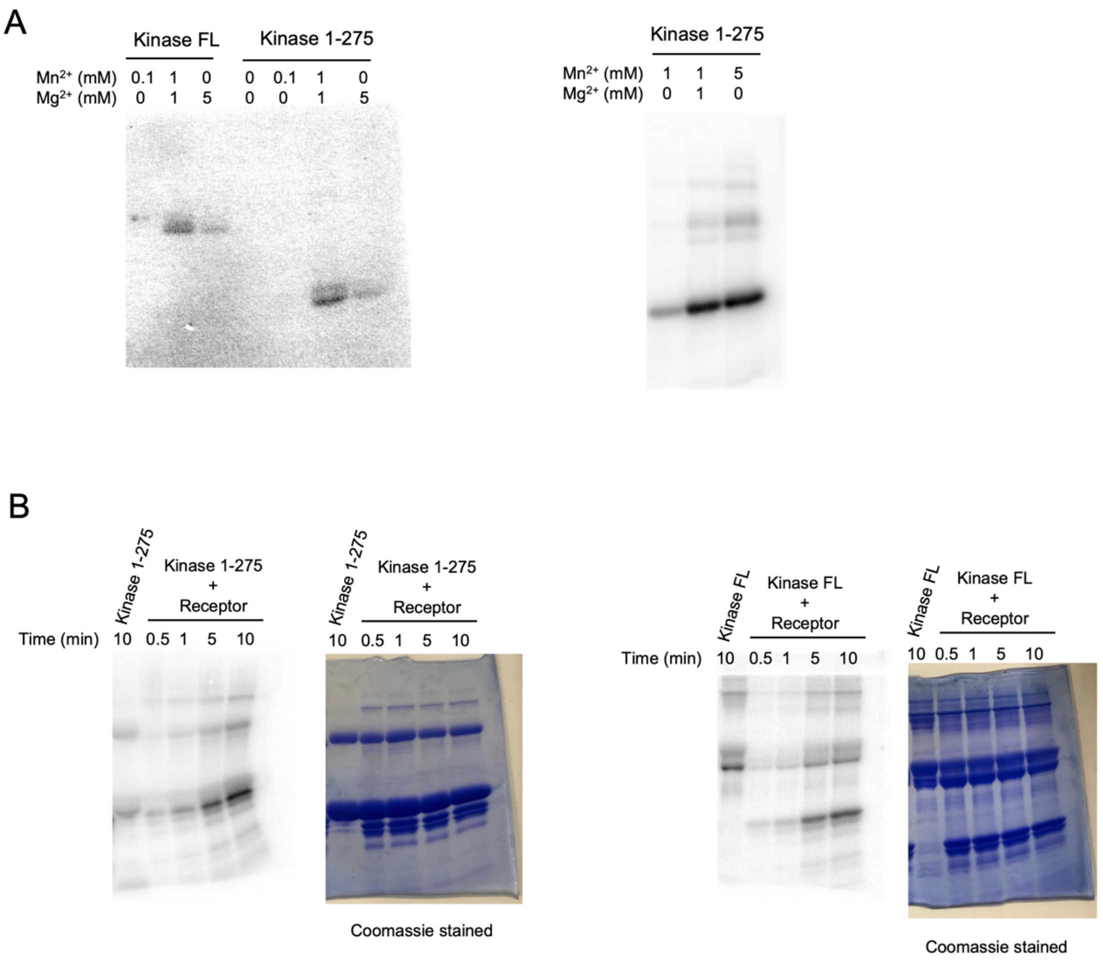

Fig. S8: Autoradiograms of kinase autophosphorylation (A) and time course of phosphorylation of the receptor (B)

A. Kinase autophosphorylation increases in the presence of $Mn^{2+}$. Thus, the kinase is $Mn^{2+}$-dependent, similar to PknB.

B. Phosphorylation of the receptor by its cognate kinase increases over a 10 minute time-span.

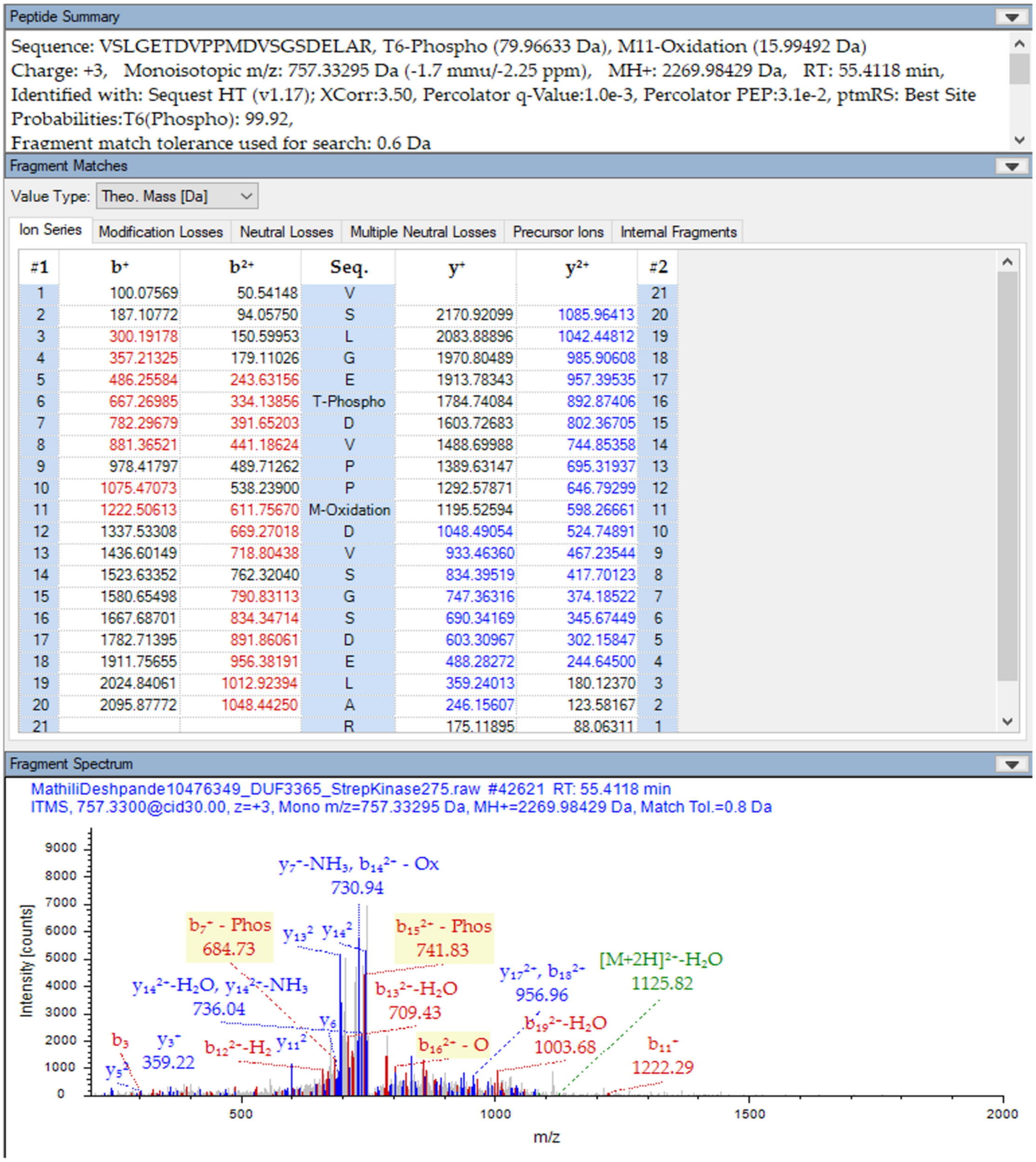

| #1 | b⁺ | b²⁺ | Seq. | y⁺ | y²⁺ | #2 |
|---|---|---|---|---|---|---|
| 1 | 100.07569 | 50.54148 | V | | | 21 |
| 2 | 187.10772 | 94.05750 | S | 2170.92099 | 1085.96413 | 20 |
| 3 | 300.19178 | 150.59953 | L | 2083.88896 | 1042.44812 | 19 |
| 4 | 357.21325 | 179.11026 | G | 1970.80489 | 985.90608 | 18 |
| 5 | 486.25584 | 243.63156 | E | 1913.78343 | 957.39535 | 17 |
| 6 | 667.26985 | 334.13856 | T-Phospho | 1784.74084 | 892.87406 | 16 |
| 7 | 782.29679 | 391.65203 | D | 1603.72683 | 802.36705 | 15 |
| 8 | 881.36521 | 441.18624 | V | 1488.69988 | 744.85358 | 14 |
| 9 | 978.41797 | 489.71262 | P | 1389.63147 | 695.31937 | 13 |
| 10 | 1075.47073 | 538.23900 | P | 1292.57871 | 646.79299 | 12 |
| 11 | 1222.50613 | 611.75670 | M-Oxidation | 1195.52594 | 598.26661 | 11 |
| 12 | 1337.53308 | 669.27018 | D | 1048.49054 | 524.74891 | 10 |
| 13 | 1436.60149 | 718.80438 | V | 933.46360 | 467.23544 | 9 |
| 14 | 1523.63352 | 762.32040 | S | 834.39519 | 417.70123 | 8 |
| 15 | 1580.65498 | 790.83113 | G | 747.36316 | 374.18522 | 7 |
| 16 | 1667.68701 | 834.34714 | S | 690.34169 | 345.67449 | 6 |
| 17 | 1782.71395 | 891.86061 | D | 603.30967 | 302.15847 | 5 |
| 18 | 1911.75655 | 956.38191 | E | 488.28272 | 244.64500 | 4 |
| 19 | 2024.84061 | 1012.92394 | L | 359.24013 | 180.12370 | 3 |
| 20 | 2095.87772 | 1048.44250 | A | 246.15607 | 123.58167 | 2 |
| 21 | | | R | 175.11895 | 88.06311 | 1 |

Fig. S9: Thr 256 in the HAMP domain of the receptor is phosphorylated by the kinase MS/MS fragmentation spectra shows that T256 is phosphorylated by the kinase.

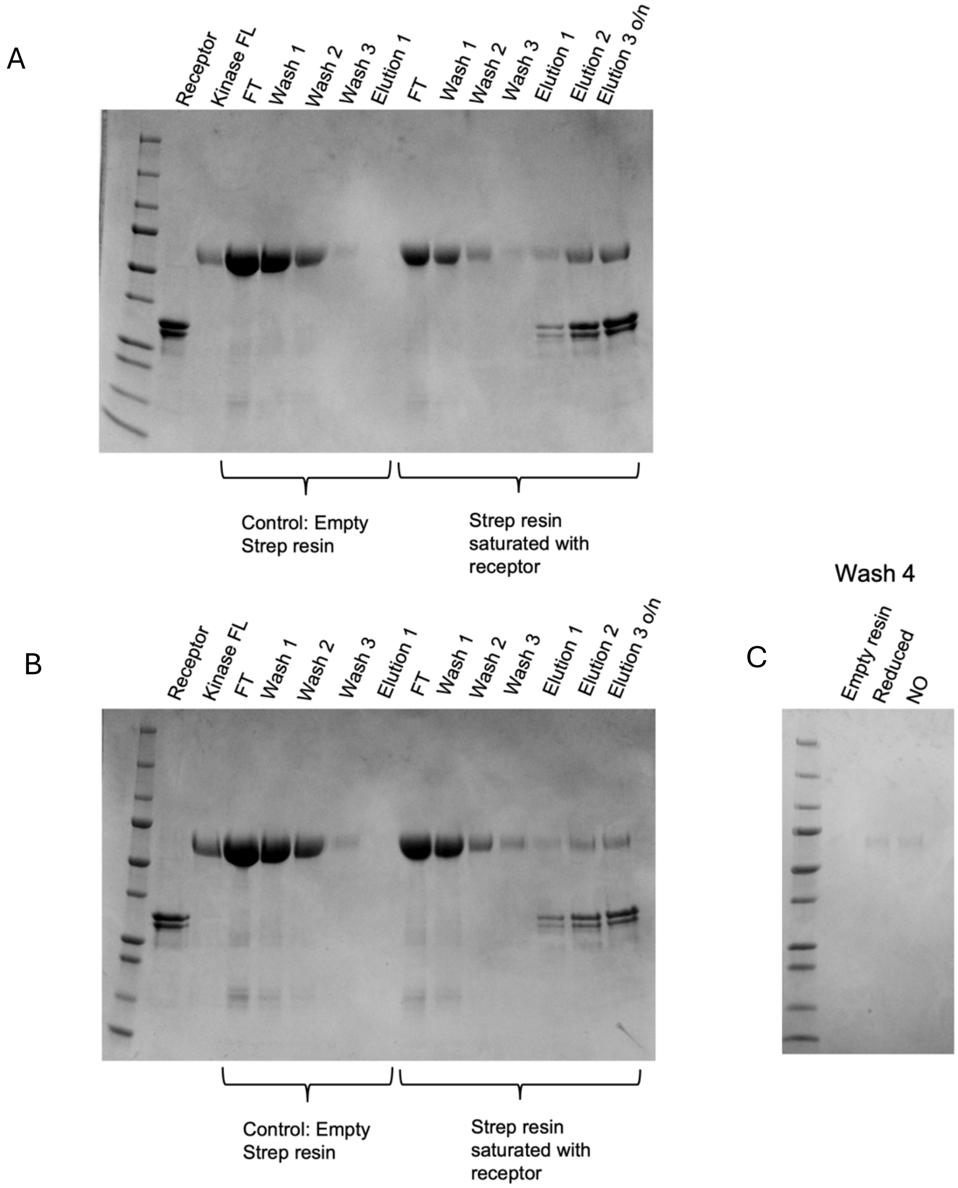

Fig. S10: Pulldown of full-length kinase by (A) ferrous (Fe(II)) receptor and (B) NO-bound receptor. Experiments were performed anaerobically. Wash 4 of empty resin, ferrous (reduced) receptor bound to resin, and NO-bound receptor bound to resin is shown in a separate SDS-PAGE gel shown in (C)

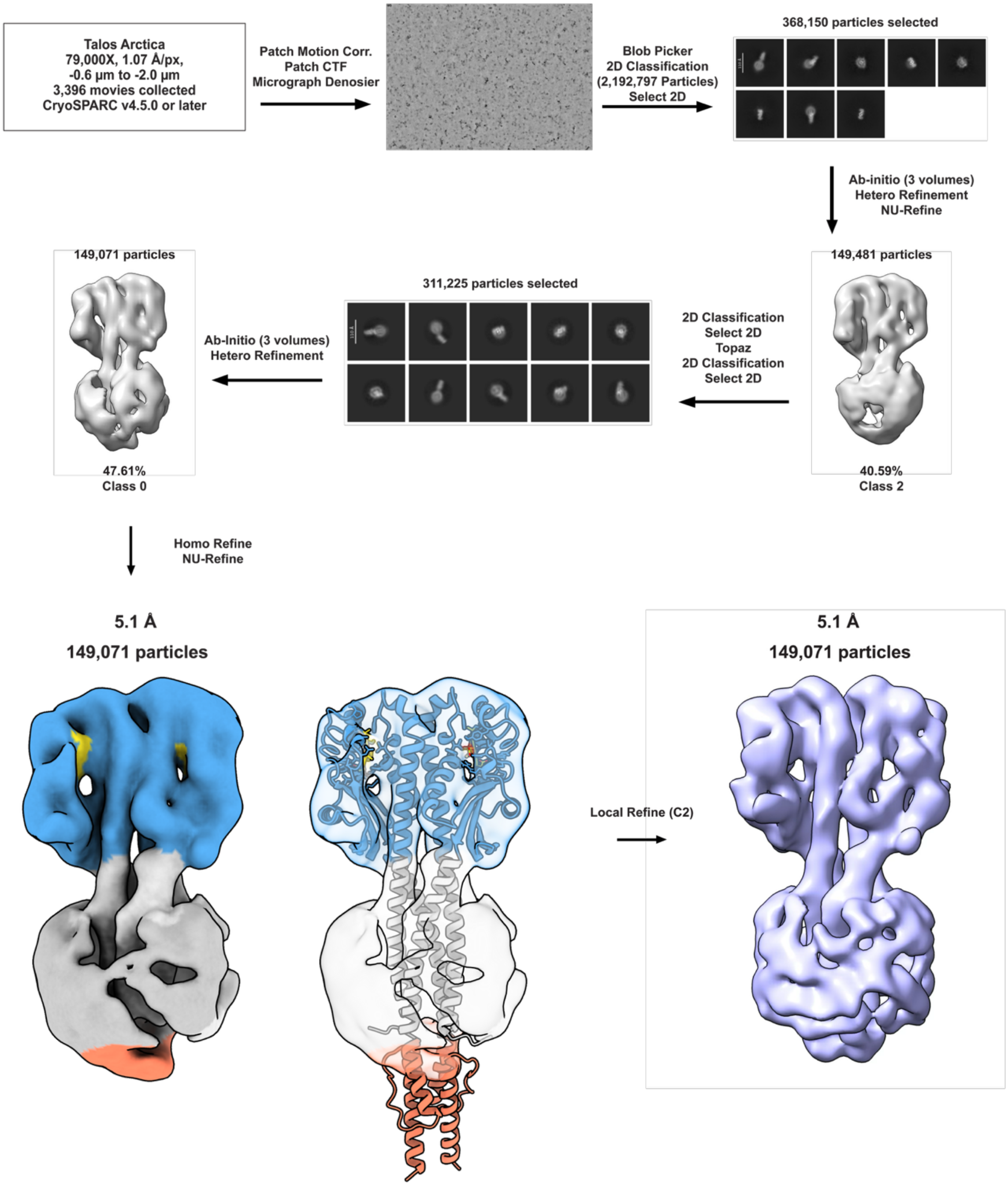

Fig. S11: Cryo-EM workflow for the receptor alone. NU – non uniform refinement.

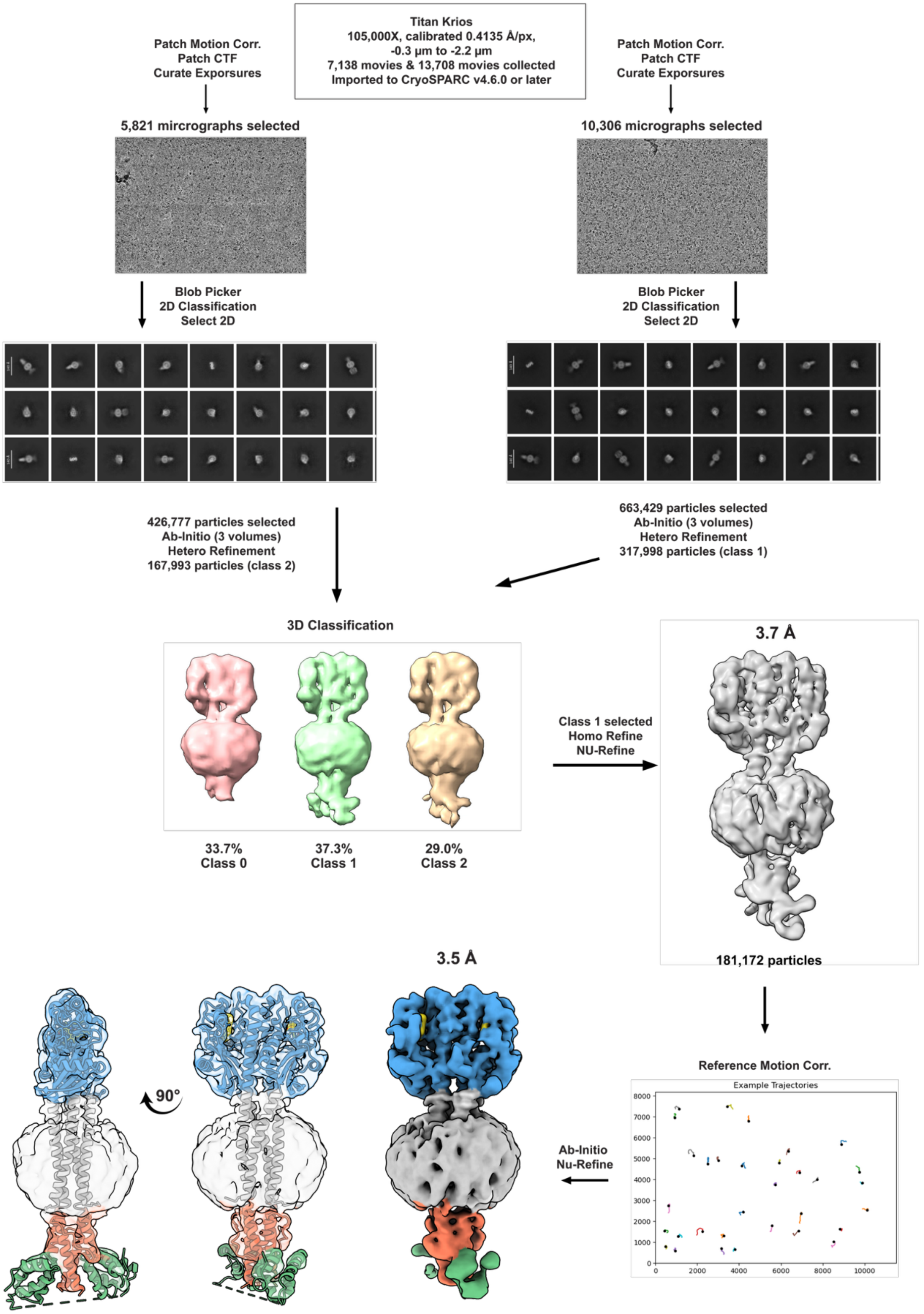

Fig. S12: Cryo-EM workflow for the receptor-kinase complex. NU – non-uniform refinement.
